# Reboot: a straightforward approach to identify genes and splicing isoforms associated with cancer patient prognosis

**DOI:** 10.1101/2020.08.18.255752

**Authors:** Felipe R. C. dos Santos, Gabriela D. A. Guardia, Filipe F. dos Santos, Pedro A. F. Galante

## Abstract

Nowadays, the massive amount of data generated by modern sequencing technologies provides an unprecedented opportunity to find genes associated with cancer patient prognosis, connecting basic and translational research. However, treating high dimensionality of gene expression data and integrating it with clinical variables are major challenges to carry out these analyses. Here, we present Reboot, an original and efficient algorithm to find genes and splicing isoforms associated with cancer patient survival, disease progression, or other clinical endpoints. Reboot innovates by using a multivariate strategy with penalized Cox regression (LASSO method) combined with a bootstrap approach, in addition to statistical tests for supporting the findings, which are automatically plotted. Applying Reboot on data from 154 glioblastoma patients, we identified a three-gene signature (IKBIP, OSMR, PODNL1) whose increased derived risk score was significantly associated with worse patients’ prognosis, even in conjunction with other well-established clinical parameters. Similarly, Reboot was able to find a seven-splicing isoforms signature (CENPF-201; MLKL-202; NUP54-201; MCF2L-201; TFDP1-207; BBS1-206; HTT-202) related to worse overall survival in 177 pancreatic adenocarcinoma patients with elevated risk scores after uni- and multivariate analyses. In summary, Reboot is an efficient, intuitive, and straightforward way for finding genes or splicing isoforms (transcripts) signatures relevant to patient prognosis, which can democratize this kind of analysis and shed light on still under-investigated sets of cancer-related genes. Reboot effectively runs on either servers or personal computers and it is freely available at github.com/galantelab/reboot.

## INTRODUCTION

The improvement of prognostic prediction and the identification of potential therapeutic targets are major interests in oncology (Hussein et al. 2018; Schirrmacher 2019). To achieve these goals, large consortiums (International Cancer Genome Consortium et al. 2010; Grossman et al. 2016) have been created, generated and made available an unprecedented amount of data, which includes clinical (*e*.*g*., survival time, tumor recurrence, treatment) and molecular information (*e*.*g*., mutation and gene expression profiles) from cancer patients and their samples. In particular, a number of studies have shown that alterations in gene expression (Huang et al. 2013; Sturtz et al. 2014; Gutierrez et al. 2019) and in splicing profiles (Yu et al. 2020; Meng et al. 2020; Liu et al. 2020) are pivotal to tumorigenesis. Once these alterations are established, researchers are often interested in pinpointing genes or splicing isoforms impacting in the prognosis of patients, which are naturally suitable therapeutic targets.

In this scenario, Cox regression models are the gold standard methodology to find genes or splicing isoforms associated with cancer patient survival. Most commonly, analyses performed on datasets with a large number of covariates are either based on simple univariate regression models or their derived forms for variable selection (Zhang 2016). However, multivariate regression models are more suitable for multifactorial phenomena due to their ability to provide synergistic and antagonistic interrelation for explanatory variables (Xing et al. 2017; Glover et al. 2015; Martinez and Kolodner 2010), a typical condition when dealing with complex diseases like cancer.

Nevertheless, such traditional models are susceptible to data idiosyncrasy. For instance, considering the high number of covariates usually present in gene expression data, it may be a challenging task to build Cox models accounting for all of them with high accuracy (Xu 2012). In a first attempt to overcome this limitation, some methods such as the Least Absolute Shrinkage and Selection Operator (LASSO) have been implemented to simultaneously estimate coefficients and treat data high dimensionality using variable selection techniques (Tibshirani 1997). Nonetheless, these implementations ordinarily exhibit poor performance for large datasets (*e*.*g*., gene expression data generated by RNA sequencing methodologies), leading to struggling in the algorithms’ convergence steps. Additionally, high collinearity and low variance of gene expression may result in incorrect estimations of the individual contributions of genes and even the identification of redundant variables in a derived model (Lesaffre and Marx 1993). Moreover, finding and testing the prognostic value or biomarker potential of a gene set is a demanding task for researchers and clinicians without extensive bioinformatics training (Zhang et al. 2019). To aid, several computational tools have been created, but still with flaws inherent to them, namely: (i) finding genes that are suitable for accomplishing the user’s goals; (ii) difficulties to determine the exact data type and even the appropriate method for user’s experiments; (iii) impossibility to customize analyses and inputs, among others (Gill et al. 2016). An easy-to-use command-line tool is routinely a worthy and more powerful option.

Here, we present Reboot, an algorithm to identify and validate gene or transcript signatures highly associated with patient prognosis. Reboot innovates by using a multivariate strategy with penalized Cox regression (LASSO method (Tibshirani 1997)) combined with a bootstrap approach (Efron 1992). Our algorithm deals with collinear variables inherent in gene expression data preventing redundancies and incorrect estimates, also/thereby removing genes with low impact on survival (*i*.*e*., low expression variance among individuals). Reboot provides complementary statistical tests to bolster gene signatures associated with patient survival or any other endpoint chosen. Furthermore, Reboot generates supporting figures, such as Kaplan-Meier curves (Jager et al. 2008) and Forest Plots (Lewis and Clarke 2001) to facilitate the interpretation of survival outcomes. Finally, Reboot seeks not only genes but also splicing isoforms (transcripts) associated with patient prognosis, successfully managing to cope with the escalation of the analysis and incorporating a deeper level of transcriptomic data interpretation to survival analyses in a practical way.

## RESULTS

### Implementation and availability

Reboot is written in R (v3.6) and comprises two major modules: regression and survival **(Figure 1)**. These two modules were designed to work independently, allowing users to identify genetic signatures using the “regression” module, and to test the significance of these signatures in prognosis using the “survival” module, possibly with additional validation datasets. Moreover, we also provide a “complete” mode option which enables the integrated execution of the two modules in case the same dataset is intended to be used in both analyses.

**Figure 1.**
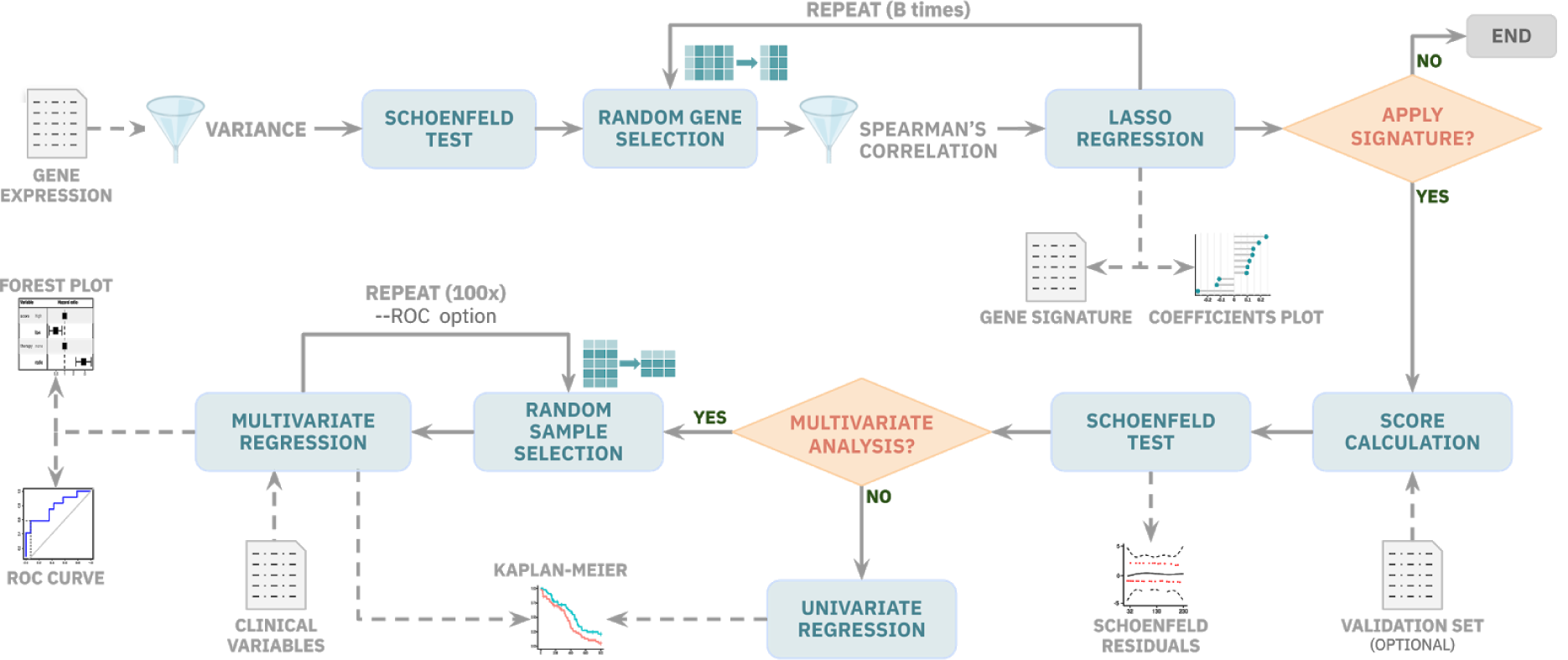
Reboot pipeline automatically integrates robust statistical tests, provides plots, and allows users to control parameters straightforwardly. In module I, gene or transcript expression data are filtered for variance and cox proportional hazard assumptions. Then, genes go through a bootstrap resampling random selection for LASSO regression and signature generation if they are not significantly correlated. In module II, a signature-based score is created and applied in survival analysis. Users are able to perform multivariate analyses, with or without bootstrap resampling and ROC curves, if clinical data are available. Plots are automatically yielded to the users.

The Reboot “regression” module is an easy-to-run step, which encapsulates statistical models to identify genes or splicing isoforms (transcripts) signatures. In brief, this module starts by checking and removing data attributes with variance lower than a user-defined or default cutoff. Next, a Schoenfeld test (Abeysekera and Sooriyarachchi 2009) is applied in a univariate way for each remaining attribute in the dataset using the packages “survival” (Therneau 2015) and “survminer” (Kassambara et al. 2019). Every attribute that fails this proportional hazard assumption test is automatically removed from the analysis. After that, a Spearman’s correlation (Akoglu 2018) filter is applied to every iteration of the bootstrap process (Efron 1992) based on the settable fraction of pairs with a correlation coefficient higher than 0.8 and a p-value < 0.05. Lastly, a bootstrap process is executed, which consists of a random sampling of attributes to be evaluated in a multivariate analysis. Regression itself is performed using a Least Absolute Shrinkage and Selection Operator (LASSO) algorithm (Tibshirani 1997) from the R packages “penalized” (Goeman 2010) and “survival” (Therneau 2015).

The next step in Reboot is the “survival” module, which is also easily executable. It receives and tests a gene/transcript signature produced in the previous (regression) module. In this step, the Reboot algorithm first produces and applies a score for all samples based on the gene/transcript signature coefficients obtained from the “regression” module and their corresponding expression values using the following formula: 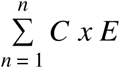, where “C” is the coefficient and “E” is the expression value. Next, the Schoenfeld test is applied to verify whether the score addresses the Cox model assumptions. Based on the median value (default) of the obtained scores, all individuals being tested are stratified into two groups, “low” or “high” score. The log-rank test (Bland and Altman 2004) is then applied in order to assess the relevance of the observed differences and to evaluate the relevance of the signature score as a prognostic factor for a given event, such as overall survival, progression-or recurrence-free survival. Next, a Kaplan-Meier survival curve is generated (Jager et al. 2008)(Schröder et al. 2011)(Jager et al. 2008) Of note, Reboot offers a multivariate option that allows extension of the survival model by additional clinical variables (*e*.*g*., therapy, age, and gender) (van Dijk et al. 2008). If this option is chosen, after applying the Schoenfeld test to all variables, multiple univariate analyses are performed and only variables under a minimal threshold (see Methods) are selected for the final multivariate model (Bradburn et al. 2003b) and illustrated in a forest plot using the R package “forestmodel” (Kennedy 2018).

Moreover, Reboot has an alternative to the use of the median value as a cutoff to stratify patients (e.g., into “low” and “high” groups, based on gene expression); this cutoff value may be defined using a Receiver Operator Characteristic (ROC) curve (López-Ratón et al. 2014)(Obuchowski and Bullen 2018)(López-Ratón et al. 2014) with the Nearest Neighbour Estimate (NNE) method and the Youden statistics (Hajian-Tilaki 2013). In this case, a patient-oriented bootstrap resampling strategy is performed using the R package “sjstats” (Lüdecke 2020). In order to derive highly confident and robust results, additional filters are applied such as null data removal, the minimum number of co-variables available and proportionality requirements (Clark et al. 2003). As a consequence, these filters ensure that the final dataset is composed of at least 70% of patients’ data present in the original one. Additionally, the final dataset als has a minimum of two co-variables to be tested with the score, whose less abundant categories’ frequencies are not smaller than 20%. After 100 cycles, the relevance frequency of each co-variable with the event is calculated and only the ones with at least 25% are plotted.

### Usage and performance

Reboot was designed to be easy-to-install and of straightforward use. To generate a genetic signature, Reboot only requires a matrix of survival data along with gene/transcript expression values as input in the form of a “.tsv” file. This file should contain as its first three columns the sample identifiers, the survival status of individuals (*e*.*g*., 0 = alive or 1 = dead) and the follow-up time, as well as gene expression values for multiple genes or transcripts across all individuals (file rows). In order to test a genetic signature, Reboot requires in addition to survival and expression data, a signature matrix with the previously produced regression coefficients (“.tsv” file), which is not necessary when “Complete” mode is run. In case a multivariate survival analysis is requested by the user, an additional file containing clinical variables to be tested should also be provided (“.tsv” file; (Bradburn et al. 2003a)). Figure S6 shows summarized examples of inputs to Reboot.

As output, Reboot generates two main textual results (“.tsv” files): (i) a list of genes or transcripts that comprise the genetic signature and their corresponding regression coefficients, which explain the contribution of each gene or transcript to the signature; and (ii) the survival impact of the signature score, including hazard ratio estimates, log-rank p-values, number of samples and median survival per group, among others. In addition, multiple plots are produced: (i) a lollipop plot, displaying the ten most significant gene coefficients comprising the signature; (ii) a coefficient histogram, displaying the distribution of all gene coefficients in the signature; (iii) a proportional hazard assumptions plot (Schoenfeld test); and (iv) a Kaplan-Meier plot **(Figure 1)**. Should the multivariate option be chosen, Reboot returns all files and plots generated in the univariate analysis plus an additional file (in “.tsv” format as well) containing the survival results of the signature score along with all other clinical variables, also visible as a forest plot. Furthermore, if the score stratification is performed with the ROC method, the curve is also plotted. Finally, a histogram of co-variable frequencies is also provided in case the multivariate option was done with bootstrap resampling.

In order to analyze the performance and features of Reboot, we built a toy dataset containing clinical and random gene expression data from The Cancer Genome Atlas (TCGA; (Grossman et al. 2016)) **Tables S1 and S2**. Correlation between the number of iterations and execution time was assessed by varying the number of iterations and keeping group size and number of instances (patients) constant in two independent tests using either a server or a laptop (see Methods for details). As expected, a linear behavior for running time was observed and server performance was slightly better than laptop. Considering Reboot modules separately, regression massively accounts for the total running time **(Figure 2A and Figure 2B)**. In another test, group size variation was also evaluated, while the number of iterations and patients was kept constant. In contrast to the previous experiment, group size variation presented a weak linear correlation. Nevertheless, the server again performed slightly better both in regression and survival modules, and time proportion between them showed a preponderance of the regression module, as expected **(Figure S2A, Figure S2B)**. Furthermore, the number of patients also presented a linear behavior when tested in an isolated way **(Figure S2C)**.

**Figure 2:**
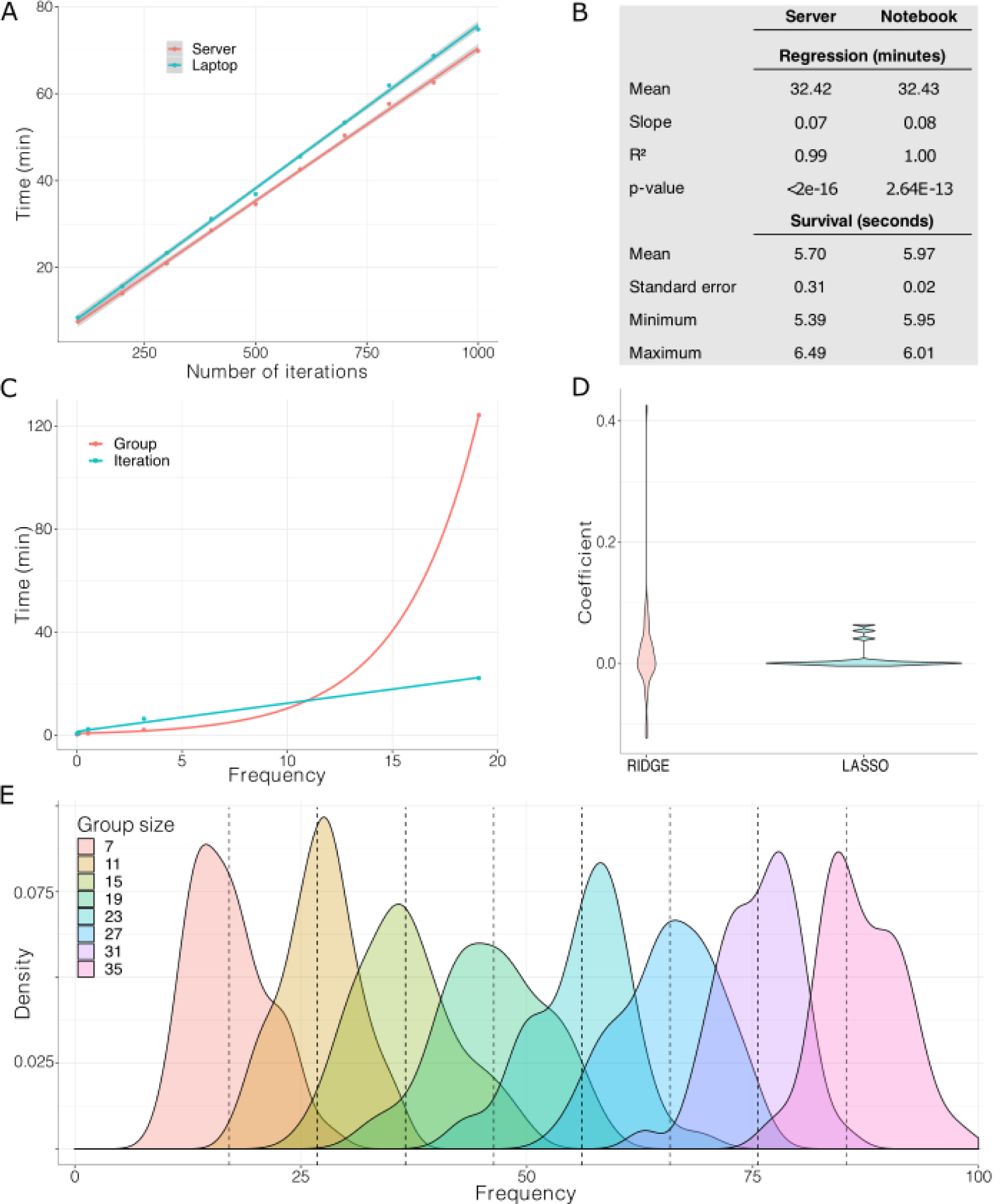
Reboot is computationally efficient. (A) Evaluation of total execution time for a Reboot complete run in a server and a laptop according to the number of iterations. Number of iterations varied from 100 to 1000 in steps of 100, keeping group size 20, 145 patients, and 50 randomly chosen genes. Survival was performed in multivariate mode. (B) Table with extracted parameters obtained in A. (C) Time assay comparing the impact of group size and the number of iterations on time of execution. Group size and the number of iterations varied from 3 to 243 in powers of 3 and from 2 to 32 in powers of 2, respectively, in both curves. Legend attribution corresponds to the variable that varied in powers of 3. (D) Coefficient distribution profile obtained from LASSO and Ridge algorithms. (E) Frequency distribution of attribute selection performed with group size variation. Theoretical average is shown in dashed lines.

The frequency of sampling for the analysis follows a distribution in which the expected average is given by equation 1, where “G” is the group size, “B” the number of iterations, and “N” the total number of attributes.

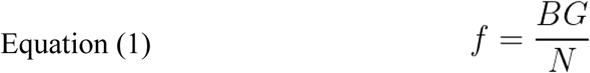

Since “B” and “G” are both directly proportional to attribute frequency, we also sought to compute time correlation of different increasing rates of group size and number of iterations with time. For this analysis, a larger group of 500 genes was randomly selected, similar to data retrieved previously **(Table S3)**. Both variables were increased by powers of 2 and 3 and multiplied, resulting in two curves containing points with the same frequency **(Figure 2C)**. Group size increase was more efficient (lower time consumption) for small frequency values when compared to the number of iterations. However, group size exhibits exponential behavior, whereas number of iterations remains linear, even for high values, indicating that increasing the number of iterations is more efficient for high attribute coverage **(Figure 2C)**. Besides time performance, large group sizes may introduce convergence issues as dimensionality increases. Reboot deals with convergence failure by reshuffling and recomputing coefficients, which may also introduce bias for large groups.

Additionally, we performed assays to compare the impact of algorithm differences and to justify why we picked LASSO instead of others. Firstly, LASSO and Ridge regressions (Hoerl and Kennard 1970) were run with a group size of 10 and 1000 iterations **(Figure 2D)**. Non-zero coefficients were extracted and a distribution was built for each analysis. As expected, the LASSO algorithm used in Reboot compresses coefficients more efficiently, denoted by the highly populated regions around 0 in relation to Ridge **(Figure 2D)**. Remarkably, LASSO also tends to keep significant coefficients closer to 0, while Ridge allows drastic distancing from coefficients **(Figure 2D)**. LASSO also exhibits its variable selection feature, compressing coefficients to zero, and leaving a lower number of significant coefficients **(Figure S1D)**.

Finally, we scrutinized the total sampling frequency of the attributes. Data obtained for **Figure S1A** was used to compute gene frequency, according to equation 1, by varying “G” **(Figure 2E)**. Mean standard deviation for all eight distributions was 4.93, contributing to a reliable uniformity of variable assessment despite the stochastic process associated with the iterative process. Obtaining a satisfactory frequency depends on the total number of attributes “N”, the group size chosen “G” and the number of iterations “B”. It is recommended that the frequency of each attribute be N/G. In accordance with equation 1, “B” may be chosen in terms of equation 2.

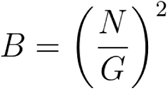

Given that a free variation of “B” performs better in terms of computational time and prevents bias, “G” may be chosen for restricted lower values and “B” estimated, with no restrictions.

### Using Reboot to identify genes related to prognosis in glioblastoma

To show how straightforward, useful and fast Reboot can be, we have applied it to a set of 1,013 protein-coding genes up-regulated in glioblastoma (GBM) in comparison to low-grade glioma (LGG) patients (log2FoldChange ≥ 2 and FDR adjusted p-value < 0.05; **Table S4**). Reboot was executed using the “regression” module parameters “-G 10 -P 0.3 -V 0.01 -B 1000” and its execution took 1.15 hours in a standard server (see Methods). As a result, we identified 255 genes associated with patients’ overall survival **(Table S5)**.

To determine whether these 255 genes could be important in GBM patient prognosis, we further investigated them. First, we performed functional enrichment analysis that revealed 131 genes (51.37%) associated with several hallmarks of cancer according to the Molecular Signatures Database (FDR < 0.01, hypergeometric test; **Table S6, Figure 3A**). Among the top 10 enriched hallmarks, we found 49 genes linked to at least 2 hallmarks relevant for glioblastoma progression and invasion, including those defining epithelial-mesenchymal transition (Iwadate 2016), encoding components of blood coagulation (Navone et al. 2019), as well as genes up-regulated in response to hypoxia (Monteiro et al. 2017) and/or by KRAS activation (Holmen and Williams 2005), among others. Genes associated with GBM patients’ survival were also enriched in a number of GO biological processes and glioblastoma-related KEGG pathways (FDR < 0.01, hypergeometric test; **Tables S7-S8, Figure 3B**). GO groups include, but are not limited to, processes related to inflammatory response, cell adhesion, proliferation, and motility, while the glioblastoma-related KEGG pathways with the greatest number of genes were proteoglycans/pathways in cancer, PI3K-Akt signaling pathway, and focal adhesion.

**Figure 3.**
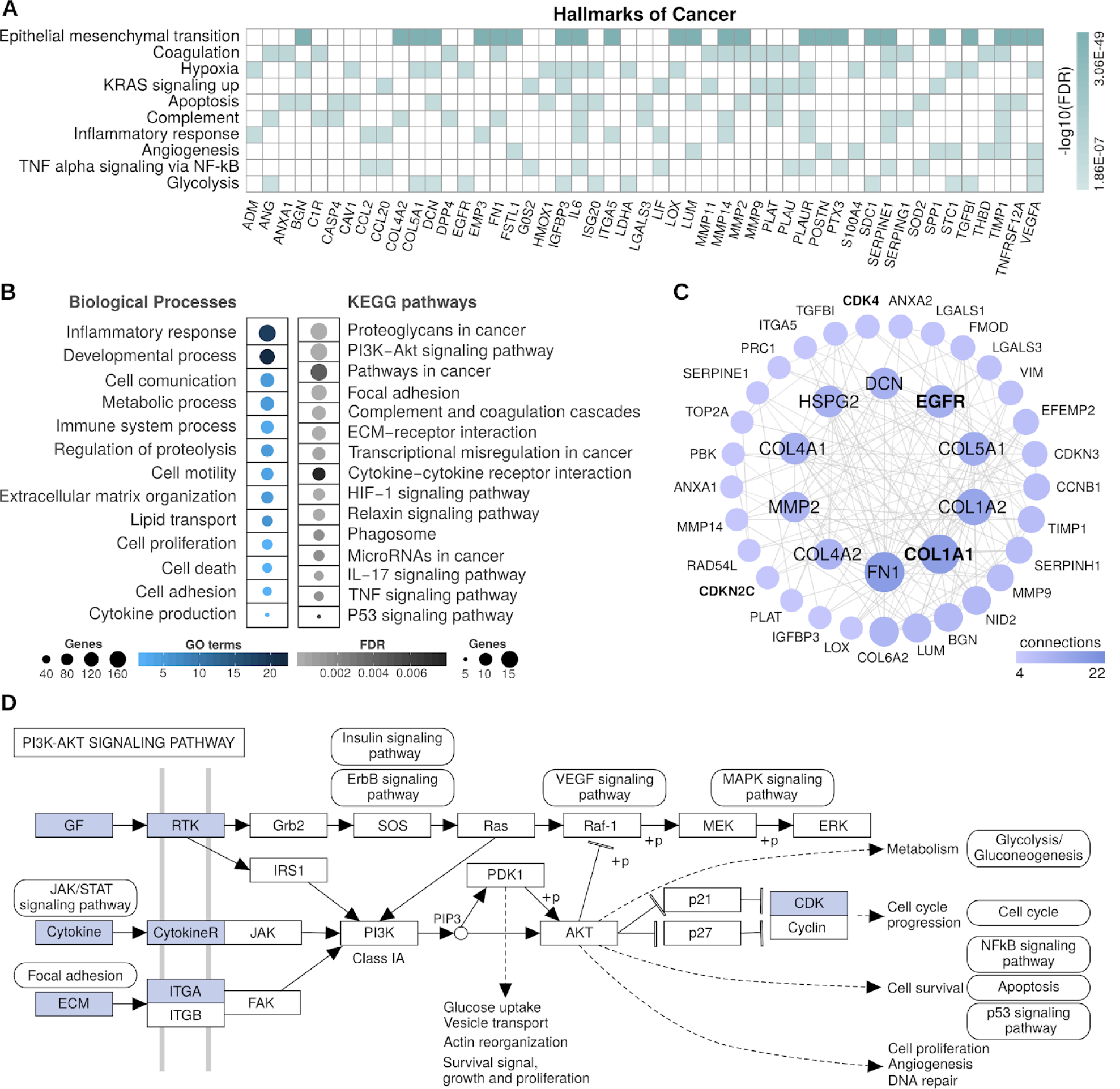
Functional enrichment analysis of genes associated with glioblastoma patients’ overall survival obtained using Reboot regression. (A) Top 10 enriched hallmarks of cancer and genes associated with at least 2 of them; (B) Groups of enriched GO biological processes and glioblastoma-related KEGG pathways; (C) Network-based on protein-protein interactions from STRING database with cancer driver genes highlighted in bold; (D) Schematic diagram of how up-regulation of 15 genes in glioblastoma may lead to activation of the PI3K-Akt signaling pathway (simplified KEGG representation with gene products highlighted in blue).

We also conducted a protein-protein interaction analysis using these 255 genes, which displayed a highly connected gene network comprising four cancer driver genes according to the Cancer Gene Census (CGC) database (https://cancer.sanger.ac.uk/cosmic/census; **Figure 3C**): COL1A1, EGFR, CDK4, and CDKN2C. Moreover, according to CGC, six other driver genes were observed in the produced signature, most of them having an oncogenic role (**Table S9**). Several genes associated with GBM initiation and progression were also observed in the network, including EGFR, MMP2, HSPG2, and various members of the collagen gene family (*e*.*g*., COL1A1, COL1A2, and COL5A1), which encode components of the extracellular matrix. Of note, fibronectin (FN1) was the top enriched gene in our network. An intracranial GBM xenograft model (Serres et al. 2014) showed that expression of FN1 promotes cell proliferation and resistance to ionizing irradiation, facilitates cell invasion and enhances angiogenic tumor growth. More recently, (Liao et al. 2018) provided evidence that fibronectin silencing in gliomas is associated with disruption of the PI3K-Akt signaling pathway and subsequent inhibition of cell proliferation, as well as promotion of cell apoptosis and senescence. Accordingly, we observed 15 genes highly expressed in GBM mostly encoding activators of the PI3K-Akt signaling pathway **(Figure 3D)**, which is frequently activated in GBM (approximately 90%; (Langhans et al. 2017; Li et al. 2016). Altogether, these 255 candidates contain many genes already reported as relevant to GBM origin, maintenance and progression, suggesting that Reboot consistently selected a gene list potentially related to prognosis in glioblastoma.

### Using Reboot to identify a minimal gene signature relevant to GBM survival

Next, we sought to determine the minimum gene set with the highest regression coefficients that are capable of explaining differences in overall survival (OS) of GBM patients using Reboot “survival” module in multivariate mode (run in docker with parameters “-M -C”; execution time ∼10 seconds in a standard laptop). Out of the total 255 genes associated with patients’ overall survival using Reboot **(Figure 4A; Table S4)**, we identified three candidates: IKBIP, OSMR, PODNL1. They are among the top 10 genes identified as the most relevant for the prognosis of GBM patients **(Figure 4B)** and are over-expressed in GBM samples in comparison to low-grade glioma (LGG) **(Figure 4C)**. Moreover, IKBIP, OSMR, PODNL1 combined score has a significant impact on survival of GBM patients (HR=0.48 95% CI: [0.32-0.71], p-value < 0.001; **Figure 4D**). The median OS for patients with a high score (> 0.34) was 335 days, yet for the low score group was 468 days. More importantly, the obtained risk score remained significant (HR = 0.53 95% CI: [0.33-0.86), p-value = 0.01, **Figure 4E**) even when considered together with relevant clinical parameters for GBM patients, including age at diagnosis, chromosome 19/20 co-gain, G-CIMP, *IDH1* mutation, and *MGMT* methylation status.

**Figure 4.**
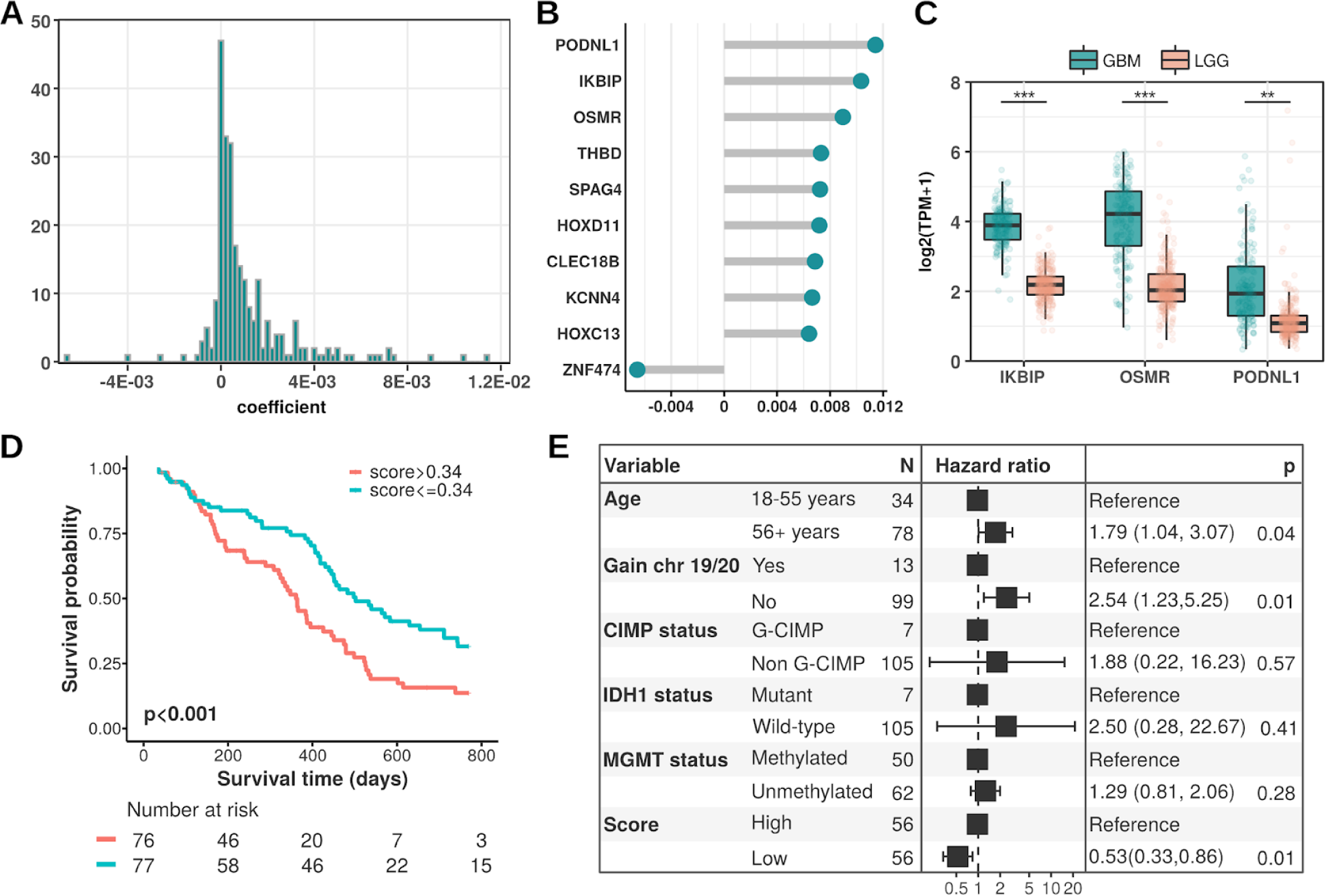
Reboot’s application on the glioblastoma dataset. (A) Histogram displaying the distribution of all gene coefficients obtained using Reboot regression; (B) Top 10 genes identified as relevant for the prognosis of GBM patients; (C) Boxplots displaying the expression values of a 3-gene signature identified in GBM patients with Reboot (Wilcoxon test; **p<0.01, ***p<0.001); (D) Kaplan-Meier curve based on the 3-gene signature score identified in GBM patients with Reboot; and (E) Forest plot of a multivariate model including the 3-gene signature score adjusted for clinical parameters relevant to prognosis in glioblastoma.

In addition, we attempted to validate this three-gene signature in an independent cohort of 71 primary glioblastoma patients from the Chinese Glioma Genome Atlas (CGGA). Similarly, higher combined scores tended to be associated with worse prognosis of GBM patients (HR=0.66 95% CI: [0.38-1.15], p-value = 0.14; **Figure S1**). The median OS for patients with higher scores (> 0.44) was 381 days versus 550 days for the low score group. Although we observed a clear separation between the higher and lower score groups in the CGGA cohort, the lack of statistical support might be explained by the smaller CGGA cohort size and sequencing depth (71 samples, ∼22.5 million *reads* on average) compared to TCGA (154 samples, ∼64.8 million *reads* on average). Thus, this result indicated that Reboot was efficient in selecting a minimal gene signature, in which a high expression is associated with worse prognosis to GBM patients.

### Finding alternative splicing isoforms signature relevant to pancreatic adenocarcinoma patients’ prognosis with Reboot

Next, we used Reboot to find splicing isoforms related to pancreatic adenocarcinoma patients’ prognosis. First, we retrieved expression data from 167 healthy pancreatic tissues from The Genotype-Tissue Expression (GTEx; (Carithers et al. 2015) and from 177 pancreatic adenocarcinomas (PAAD) samples from TCGA (Grossman et al. 2016). For cancer data, we also obtained their respective clinical information (Liu et al. 2018). Next, by using Kallisto (Bray et al. 2016) and SUPPA2 (Trincado et al. 2018) tools, we found a complete set of alternative splicing isoforms (ASI) in pancreatic adenocarcinoma, which fed the Reboot’s algorithm to perform the signature (part I) and the survival (part II) analyses (**Figure 5A)**.

**Figure 5.**
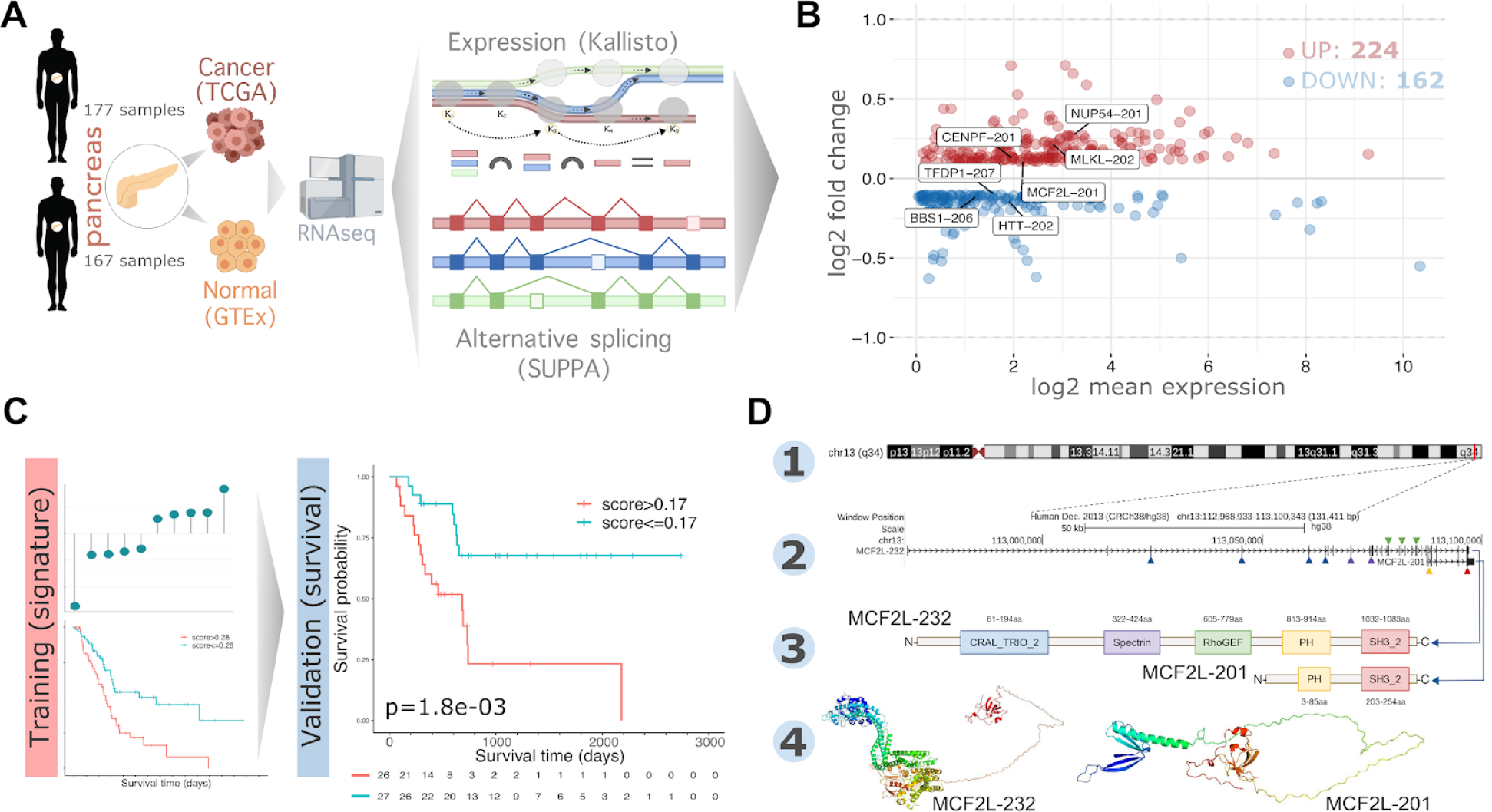
Reboot selects alternative splicing isoforms associated with pancreatic adenocarcinoma tumorigenesis and its patient’s prognosis. (A) Selection of alternative splicing isoform (ASI) based on transcript expression data from healthy (GTEx) and tumoral (TCGA) pancreas. (B) MA plot showing the mean expression (in TPM) and ΔPSI (percent spliced in) values of all ASI. Highlighted ASIs compose the seven-transcripts signature generated with Reboot. (C) ASI data was split into training (70%) and validation (30%) set to find a transcript signature in survival analysis. Kaplan-Meiers made by Reboot when using both the training (HR: 0.4428 [0.2719 - 0.7211]; p = 8e-04) and the validation dataset (HR: 0.2791 [0.1191 - 0.6541]; p = 0.0018) showed a worse survival outcome for patients with higher scores (above median value). Follow-up time (days) is shown in the bottom for each group. (D) MCF2L mapping on the reference genome (1). Canonical (longer: MCF2L-232) and ASI (shorter: MCF2L-201) isoforms, respectively(2). Protein domains encoded from canonical (MCF2L-232) and ASI (MCF2L-201) isoforms, respectively. (3). Predicted 3D protein structure for canonical (MCF2L-232) and ASI (MCF2L-201) transcripts (4).

We found 386 significantly ASI, of which 224 and 162 were up-regulated and down-regulated, respectively, in PAAD versus healthy pancreas tissue **(Figure 5B)**. To prove the Reboot robustness in candidate selection, we randomly split the initial ASI data into training (70%) and validation (30%) sets **(Figure 5C)**. When applying the “regression” module on the training dataset (parameters: -B 100 -G 10 -P 0.3 -V 0.036 -F FALSE; its execution took 4.71 minutes in a standard laptop), a signature with 62 transcripts emerged **(Table S10 and Figure S3)**. After setting a cutoff of 0.035 to coefficients, Reboot found a minimal signature of seven transcripts (CENPF-201; MLKL-202; NUP54-201; MCF2L-201; TFDP1-207; BBS1-206; HTT-202) presenting significant survival results **(Figure 5C and S4, Table S10)**.

When testing the signature with module II (survival) of Reboot on the validation dataset (53 patients; parameters: -M TRUE -R FALSE -F FALSE; execution time ∼5 seconds in a standard laptop), we found that patients with higher scores (values above the median) had worse overall survival (HR: 0.2791 [0.1191 - 0.6541]; p-value = 0.0018; **Figure 5C and Figure S4)**. The median OS for patients in the high score group (> 0.17) was 684 days, whereas this value could not be calculated in the low score group since less than half of the patients died. Furthermore, this result remained statistically significant after the multivariate analysis, accounting for relevant clinical variables such as age, gender, race, Tumor Node Metastasis (TNM) classification (Amin et al. 2017), histology, and grade (HR: 0.3806 [0.1569 - 0.923]; p-value = 0.0326; **Figure S4)**. Additionally, the same results were observed when applying the score on the training dataset, as expected **(Figure S4)**, where the median OS for patients in the high score group (> 0.28) was 517 days versus 1332 days for the low score group. Of note, other endpoints such as Disease-Specific (DSS), Progression-Free (PFI) or Disease-Free Interval (DFI) may be used instead in order to better fit the data and meet survival requirements (Liu et al. 2018).

Further investigation was performed for transcripts with major contributions to the genetic score **(Figure S4)**. MCF2L-201, which had a significant positive score, lacks three protein domains: RhoGEF, Spectrin and CRAL_TRIO_2, which are all present in the canonical isoform MCF2L-232 **(Figure 5D)**. However, MCF2L-201 maintains PH and SH3_2, which are also present in the canonical isoform. Another isoform, HTT-202, which scored negatively in our signature, lacks the huntingtin protein region DUF3652, present in the canonical isoform HTT-201 **(Figure S5)**. Taken together, these results demonstrate that Reboot is effective not only to identify relevant genes but also splicing isoforms related to cancer.

## DISCUSSION

In the past few years, advances in RNA sequencing technology have provided us an unprecedented opportunity to find novel gene signatures acting as prognostic or diagnostic biomarkers in cancer (Byron et al. 2016). Notwithstanding, treating high dimensionality of gene expression integrated with clinical variables are major challenges when performing this analysis, notably by researchers without extensive training in computational biology. It is therefore an urgent task to establish robust and straightforward methods capable of handling ultra-large datasets and finding these genes. Here we describe Reboot, a user-friendly algorithm to seek, evaluate, and validate genes and splicing isoforms signatures acting as prognostic or diagnostic biomarkers in cancer. Reboot is original and efficient: i) it combines a multivariate strategy with penalized Cox regression (LASSO method) and a bootstrap approach, plus a variety of statistical tests to find genes or transcripts candidates; ii) it is easy-to-use, well documented, and of simple installation; iii) it includes effortless steps to plot its results and to facilitate data interpretation and further analyses.

Availability of tools that manage to escalate genetic score analysis with high dimensional datasets, such as those found in gene expression data, are scarce. Beyond that, to the best of our knowledge, there is no state-of-the-art pipeline that integrates posterior validation, including clinical data, gene, and splicing isoforms expression. As genetic analyses get wider in order to capture the complexity of intricate diseases such as cancer (Frampton et al. 2013; Cheng et al. 2015; Campesato et al. 2015)), coverage of the transcriptome with more genes or transcripts (splicing isoforms) becomes crucial, which significantly raises the dimension of input datasets. Moreover, beyond all implemented filters that automate pre-processing, bootstrapping strategy allows a wide range of dataset dimensions. For instance, the number of iterations has a strong linear correlation with time. The association with the LASSO algorithm, which presents a strong variable selection feature, balances score attributes in the analysis, yielding less false positives but still exhibiting gains of a multivariate analysis. All of these approaches were included in Reboot, which introduces a robust, complete, and computationally efficient algorithm.

We selected and tested Reboot on data from glioblastoma (GBM) and pancreatic adenocarcinoma (PAAD), two cancer types presenting a poor survival rate and limited therapeutic options for their patients (Kleeff et al. 2016)(Di Carlo et al. 2019)(Kleeff et al. 2016). First, we identified prognostic genes in GBM, which were associated with a number of processes relevant for glioblastoma tumorigenesis, progression, and invasion (*e*.*g*., epithelial-mesenchymal transition, inflammatory response, and cell proliferation). This list includes genes already described as related to GBM or other gliomas. In particular, the epidermal growth factor receptor (EGFR) is a primary driver of glioblastoma tumorigenesis, contributing mainly to cell proliferation and invasion (Huang et al. 2009). MMP2 is also highly expressed in gliomas and it was recently associated with stimulation of vasculogenic mimicry in glioma cells (Liu et al. 2019). HSPG2, in glioma tissues, is associated with the transformation of brain extracellular matrix into tumour microenvironment and represents a negative prognostic factor in overall and relapse-free survival (Ma et al. 2018; Kazanskaya et al. 2018).

Next, by using the “survival” module in multivariate mode, Reboot found a signature containing three genes (IKBIP, OSMR, and PODNL1) associated with GBM patients’ overall survival. Interestingly, they have emerged as prominent genes in glioblastoma: i) IKBIP has been described as a novel p53 target with pro-apoptotic activity, whose high expression is associated with poor prognosis in GBM (Cao et al. 2019; Long and Li 2019). Although in our results *MGMT* methylation was not considered a significant co-variable, another study has identified gene IKBIP as part of a signature that predicts prognosis only in GBM patients with methylated *MGMT* promoter (Wang et al. 2016). OSMR, characterized as a novel key regulator of glioblastoma tumorigenesis through EGFRvIII-STAT3 signaling, also correlates with poor prognosis in GBM patients both independently and also as part of a 4-gene signature (Mohan et al. 2017; Cao et al. 2019). Interestingly, PODNL1 encodes a protein involved in extracellular matrix formation with an unclear role in GBM tumorigenesis. The latter gene up-regulation has also been correlated with the poorest survival rates in GBM patients in distinct studies (Shergalis et al. 2018; Yan et al. 2013). Altogether, Reboot identified a valuable set of genes to be further and deeper investigated in GBM.

Second, we used Reboot to seek for alternative splicing isoforms associated with pancreatic adenocarcinoma (PAAD) patients’ prognosis. As illustrated in our analyses, a genetic score obtained from differentially expressed transcripts stratifies patients with bad and good prognosis as efficiently as from gene analyses. Interestingly, a signature score with only seven transcripts was enough to yield statistical significance in the survival analysis of PAAD patients. Among them, only three isoforms are canonical (CENPF, MLKL, NUP54). Some of these genes (*e*.*g*., CENPF, MLKL, TFDP1, MCF2L) have a known influence on cancer, while others (*e*.*g*., NUP54, BBS1, and HTT) have been superficially studied in a tumoral context. CENPF, for instance, was already related to worse outcomes and survival in several cancer types (R. Li et al. 2020; H.-B. Liu, Huang, and Luo 2020; Garcés et al. 2020; Ying Liu et al. 2020). Another great example is the MLKL gene. This gene was shown to be up-regulated in pancreatic cancer, as we observed with Reboot, especially under the context of invasion (Ando et al. 2020). The transcription factor TFDP1 is another gene whose functions remain uncovered in cancer, but significant somatic copy number alterations and corresponding somatic gene expression changes were observed in papillary thyroid carcinomas (Yang et al. 2020). Additionally, it is considered a prognostic marker in liver cancer (unfavorable), stomach cancer (favorable) and renal cancer (favorable) according to The Human Protein Atlas (Uhlen et al. 2017). Inconsistencies in these results may have arisen from a possible divergence of the role of different isoforms from this gene. Our results indicate that an isoform (TFDP1-207, down-regulated in our analysis) other than the canonical (TFDP1-201, up-regulated in our data) is of great significance for PAAD patient prognosis, an evidence that more detailed scrutiny is required for this gene (https://www.proteinatlas.org/ENSG00000198176-TFDP1/pathology). Taken together, it is clear that transcript-centered analysis may shed light on more detailed molecular mechanisms that would not be possible in a gene-based analysis. Therefore, the results provided by Reboot seem trustworthy and robust.

Nucleoporin 54 (NUP54), on the other hand, has not been extensively studied in cancer. Even though an association between larger tumor size and loss of heterozygosity (LOH) in NUP54 was found in patients with hepatocellular carcinoma, potentially playing an important role in its development and progression (G.-L. Huang et al. 2010). Moreover, it was recently reported a novel function for NUP54 in the response to ionizing radiation and the maintenance of homologous recombination-mediated genome integrity in cell lines (Rodriguez-Berriguete et al. 2018).

Among the best-scored transcripts, MCF2L-201, which was found to be up-regulated in PAAD, is a compelling example. The canonical isoform of the MCF2L gene (MCF2L-232) encodes DBL from the Guanine Exchange Factor protein family, known to directly interact and regulate important factors for cell cycle such as Cdc42 and RhoA complexes (Jaiswal et al. 2013). It has been shown that the minimal and sufficient catalytic activity of DBL is composed of a DH and a PH domain linked in tandem (Cerione and Zheng 1996). Although MCF2L-201 does not present a DH domain, it keeps a PH and a SH3 domain. PH domains perform essential contact with Cdc42 and RhoA in the DBL structure (Snyder et al. 2002). They are also known to be responsible for protein subcellular localization and phosphoinositide interaction (Zheng et al. 1996; Han et al. 1998). Moreover, SH3 (Src homology 3) domains are abundant in oncogenic pathways such as cell migration and proliferation, distributed along with many different protein structures (Birge et al. 1996). SH3 domains have also been implicated in pancreatic cancer specifically due to its relevance for oncogenic pathways (Thalappilly et al. 2008). Although a few isoforms of MCF2L have been initially explored, such as MCF2L-203 - which does not catalyze guanine nucleotide exchange on CDC42 - and MCF2L-205 - which, on the other hand, activates CDC42 (Ueda et al. 2004) - MCF2L-201 requires further investigation. Details about how the PH-SH3 protein may act and its role in pancreatic cancer deserve deeper analyses, even though our study provides some guidance on that.

Despite the fact that Huntingtin is mostly known to cause Huntington disease by the expansion of the trinucleotide CAG in its first exon, it has a wide tissue expression and its trinucleotide expansion has been correlated to tumor progression, including metastasis, and inversely correlated to carcinogenesis (Thion and Humbert 2018). Huntingtin transcript HTT-202 is non-canonical and we found it down-regulated in pancreatic tumors. Its protein structure presents neither the characteristic polymorphic trinucleotide repetitive region nor the main huntingtin annotated domain: DUF3652; thus, its function is an enigma. A similar case involves the BBS1 gene since it is most known for its association with the Bardet-Biedl Syndrome (BBS) (Forsythe and Beales 2003)(Beales et al. 2000)(Forsythe and Beales 2003). More interesting is the fact that higher expression of BBS1 was related to better survival in patients with malignant pleural mesothelioma (Vavougios et al. 2015), although in our PAAD signature this gene was down-regulated. Furthermore, BBS1 was part of a 15-gene signature associated with bone metastasis in breast carcinomas. Specifically, its up-regulation was correlated to the epithelial to mesenchymal transition status of the tumor (Savci-Heijink et al. 2016). Thus, Reboot’s algorithm makes splicing isoform expression analysis feasible in cancer prognosis,also providing clues to decipher unclear molecular mechanisms of tumorigenesis and cancer progression.

## CONCLUSIONS

In conclusion, here we present Reboot, an algorithm to seek, evaluate, and validate genes and transcripts signatures acting as prognostic or diagnostic biomarkers in cancer. Reboot brings novelties by combining a multivariate strategy with penalized Cox regression (LASSO method) and bootstrap approach, plus a variety of statistical tests to find genes or transcripts candidates. Reboot shows its usefulness by identifying prognostic genes and a minimal set of genes associated with glioblastoma patients’ survival and a splicing isoforms signature associated with pancreatic adenocarcinoma. We believe that Reboot will be of immediate interest to the cancer research community because it will accelerate and democratize the search for genes and transcripts biomarkers, even by researchers and clinicians without extensive bioinformatics training.

## METHODS

### Usage and performance

Expression and clinical data from TCGA were obtained from individuals that presented only a single primary glioblastoma tumor by an *in-house* R script (toyfordocker.R found in https://galantelab.github.io/reboot). Gene expression was obtained from processed datasets in FPKM. The same 50 randomly picked genes were used in all essays with exception of concomitant group size and number of iterations variation, in which 500 genes were randomly picked. For time comparisons, laptop and server specifications are: CPU: Intel(R) Xeon(R) Silver 4114, 2.20 GHz, 128 GB of ram; and CPU: Intel(R) Core (TM) i7-8550U 1.80 GHz, 16GB of RAM, respectively. All-time essays were computed with the parameter “M” and all others were set default unless otherwise stated. All linear regressions (Pearson) and plots were generated in R.

### Gene expression profiles

Mapped RNA-seq and clinical data of 154 and 248 samples from patients with primary glioblastoma (GBM) and low grade glioma (LGG grade II), respectively, were downloaded from TCGA (Grossman et al. 2016). We also obtained RNA sequencing and clinical data of 71 samples from primary GBM patients from the Chinese Glioma Genome Atlas (CGGA) for validation purposes. All datasets were processed using Kallisto (Bray et al. 2016) with GENCODE (v29) as reference to the human transcriptome. Gene-level counts (TPM normalized) were obtained using tximport (Soneson et al. 2015).

### Differential gene expression

Differential gene expression of GBM versus LGG-II samples from TCGA was performed using DESeq2 (Love et al. 2014), and we considered as up-regulated only genes presenting a log2FoldChange ≥ 2 and false discovery rate (FDR) adjusted p-value < 0.05.

### Functional analyses

For Gene Ontology enrichment analysis, we used ShinyGO (Ge et al. 2019) and REVIGO (Supek et al. 2011) web tools. ShinyGO was also used to evaluate cancer hallmarks from the Molecular Signatures Database - MSigDB (www.gsea-msigdb.org/gsea/msigdb/) and KEGG pathways (www.genome.jp/kegg/). Only functional terms with an FDR < 0.01 (hypergeometric test) were considered relevant. Protein-protein interaction analysis was performed in Cytoscape (Shannon et al. 2003) using the STRING database (Szklarczyk et al. 2019).

### Transcript expression profiles

Mapped RNA-seq data from 177 tumor patients with pancreatic adenocarcinoma from TCGA (Grossman et al. 2016), along with their respective clinical data (J. Liu et al. 2018), were downloaded. The TCGA dataset was reprocessed using Kallisto (Bray et al. 2016) with GENCODE (v29) as reference to the human transcriptome and transcript-level counts (TPM normalized) were obtained using tximport (Soneson et al. 2015). We directly downloaded transcript-level TPM normalized data of pancreatic samples from 167 healthy individuals from The Genotype-Tissue Expression (GTEx; Carithers et al. 2015) through the UCSC Xena portal (toil.xenahubs.net/). Only transcripts available in both datasets were further considered in the analyses.

### Differential transcript expression

Analysis of differential transcripts usage between tumoral (pancreatic adenocarcinoma) and healthy pancreatic samples from TCGA and GTEx was performed using SUPPA2 (Trincado et al. 2018; version 2.3). We considered as significant only transcripts presenting a |ΔPSI| ≥ 0.1 and FDR adjusted p-value ≤ 0.05.

### 3D structure prediction

MCFL2-201, MCF2L-232, HTT-201 and HTT-202 transcript sequences were submitted to ORFfinder (Tatusov and Tatusov 2007) with default values. The longest positive open reading frames (ORFs) were then submitted to Pfam (Punta et al. 2012). Finally, the amino acid sequences of both transcripts were submitted to Phyre2 ((Kelley et al. 2015)version 2.0(Kelley et al. 2015) for 3D structure prediction in “intensive” mode.

### Availability and updates

Reboot is implemented as an R package that is freely available under the GNU General Public Licence version 3 (GPL3) at https://galantelab.github.io/reboot/. Reboot updates will be announced at its webpage and each respective Docker image will be released along with new versions.

