## Supplemental for "Reboot: a straightforward approach to identify genes and splicing isoforms associated with cancer patient prognosis"

2 - Programa Interunidades em Bioinformatica, Universidade de São Paulo, Sao Paulo, SP.

3 - Departamento de Bioquimica, Universidade de Sao Paulo, SP, Brazil.

* These authors contributed equally to this work

#### Supplementary Figures

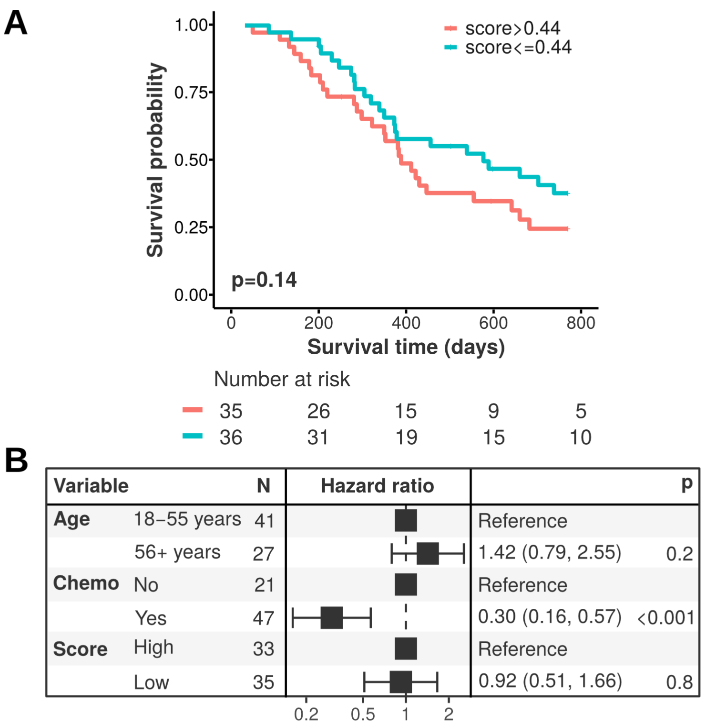


**Figure S1: Validation of the 3-gene signature score in glioblastoma patients from the CGGA cohort.** (A) Kaplan-Meier curve based on the 3-gene signature score; (B) Forest plot of a multivariate model including the 3-gene signature score along with clinical parameters relevant to prognosis in glioblastoma.

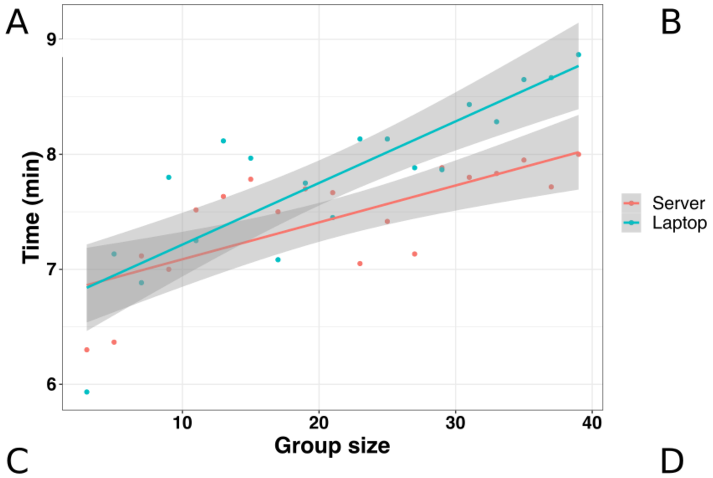


B

| | Server | Notebook |
|---|---|---|
| | **Regression (minutes)** | |
| Mean | 7.30 | 7.57 |
| Slope | 0.04 | 0.05 |
| $R^2$ | 0.55 | 0.69 |
| p-value | 7.10E-08 | 7.45E-06 |
| | **Survival (seconds)** | |
| Mean | 5.94 | 6.11 |
| Standard error | 0.22 | 0.08 |
| Minimum | 5.60 | 6.01 |
| Maximum | 6.35 | 6.24 |

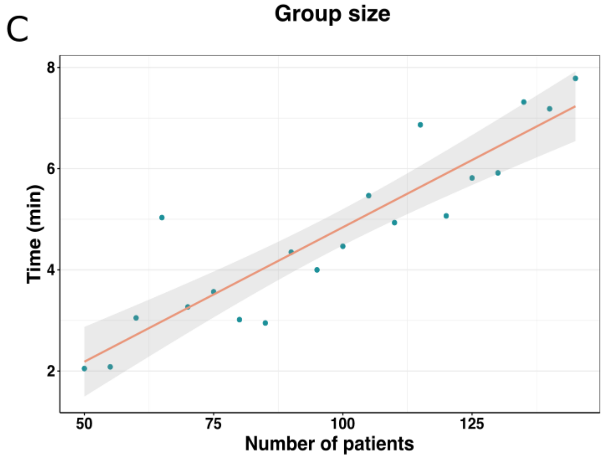


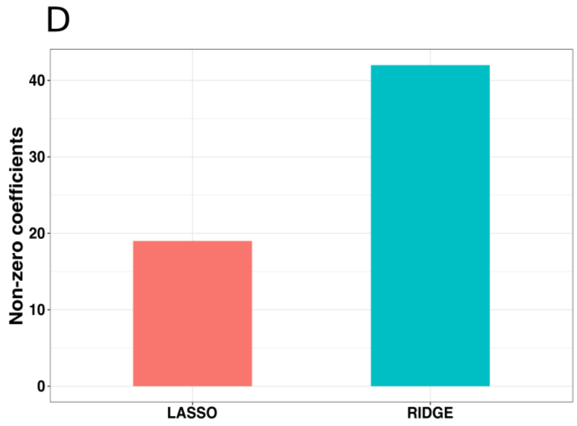


**Figure S2: Computational assessment of Reboot.** (A) Evaluation of group size impact on time performance of Reboot. Number of iterations was set to 100 and remaining parameters default. (B) Table showing numerical results for assay performed in A and split by module. (C) Run time analysis for instances (patients) variation. Processes were run in the laptop described in section "Usage and performance". Group size was set to 20, the number of iterations to 100, and remaining parameters default. (D) Number of non-zero coefficients obtained by running Ridge and LASSO algorithms with group size set to 20, number of iterations set to 500, and remaining parameters default.

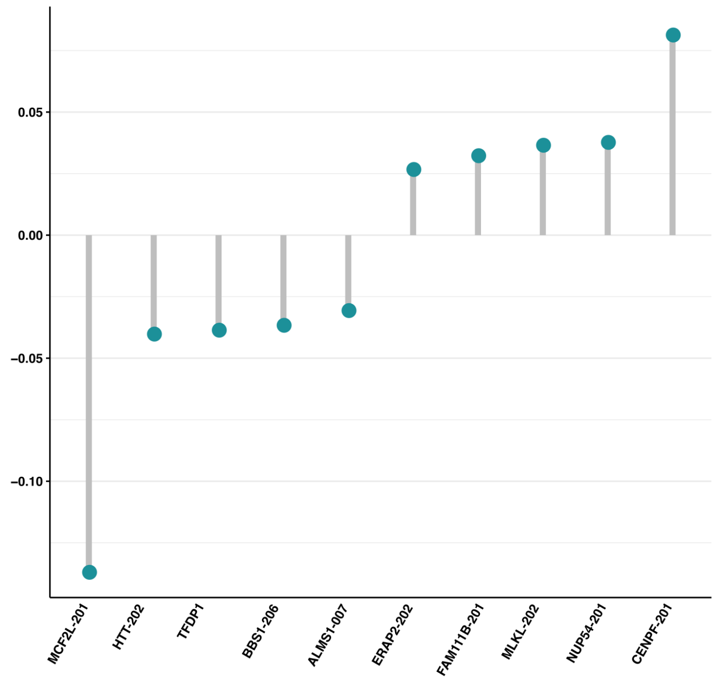


**Figure S3: Reboot's Lollipop plot.** Top 10 transcripts present in generated signature, which are relevant for PAAD survival, where the higher the coefficients (in module), the greater the influence of that transcript to the final patients' outcome.

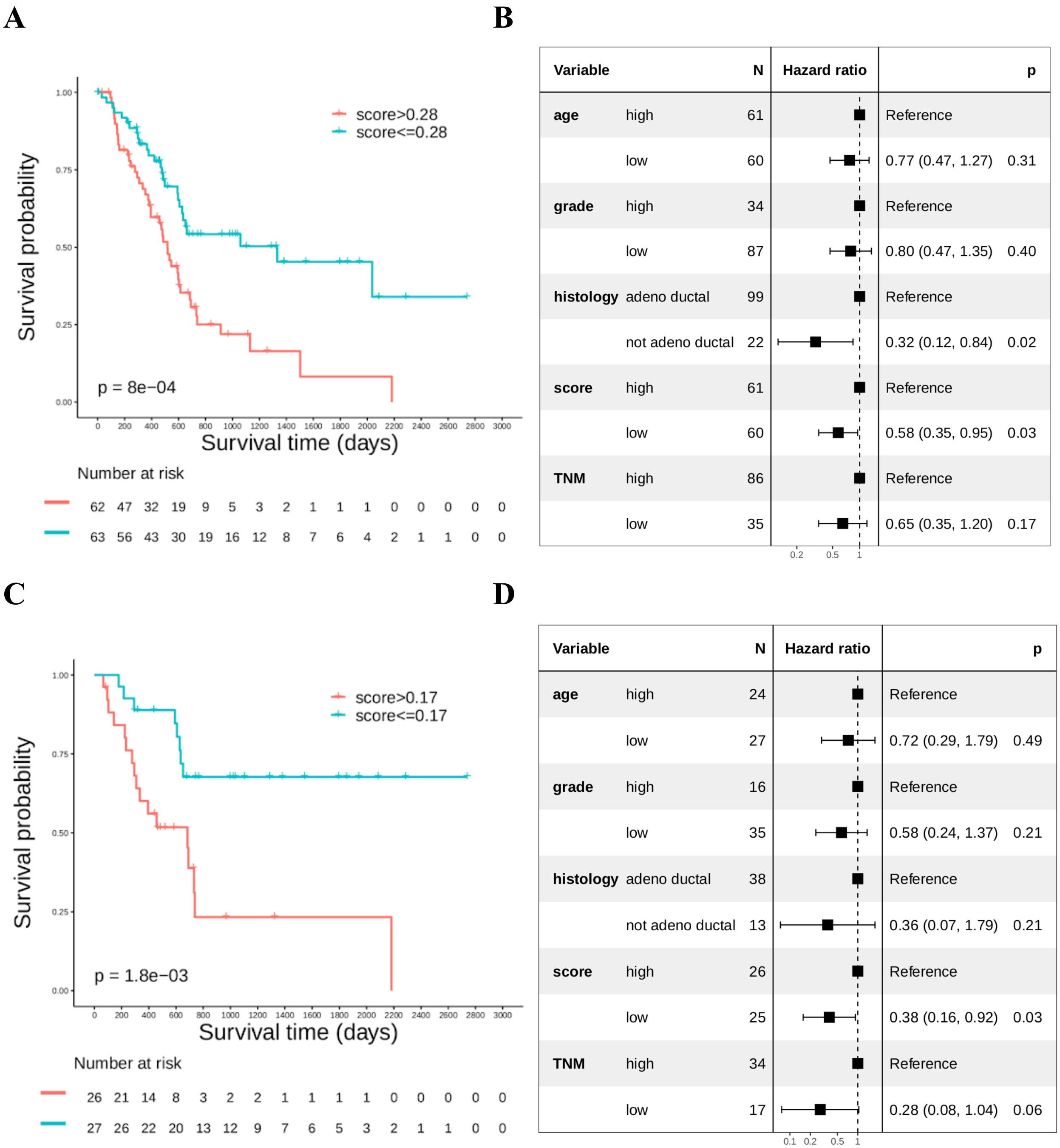


**Figure S4: Reboot's outputs of PAAD outcomes.** Univariate and multivariate survival analyses for training (A) and (B) and validation (C) and (D) datasets. (A) Kaplan-Meier displaying that patients with higher scores (above median value) have worse prognosis. Follow-up time (days) is shown below curves. (B) Forest plot showing the significance of the score in predicting patients' outcome with correction for other relevant clinical variables. Similarly, (C) Kaplan-Meier curve evidencing that patients with higher scores (above median value) have worse prognosis. Follow-up time (days) is shown below curves. (B) Forest plot confirming the significance of the score in predicting patients' outcome with correction for other relevant clinical variables.

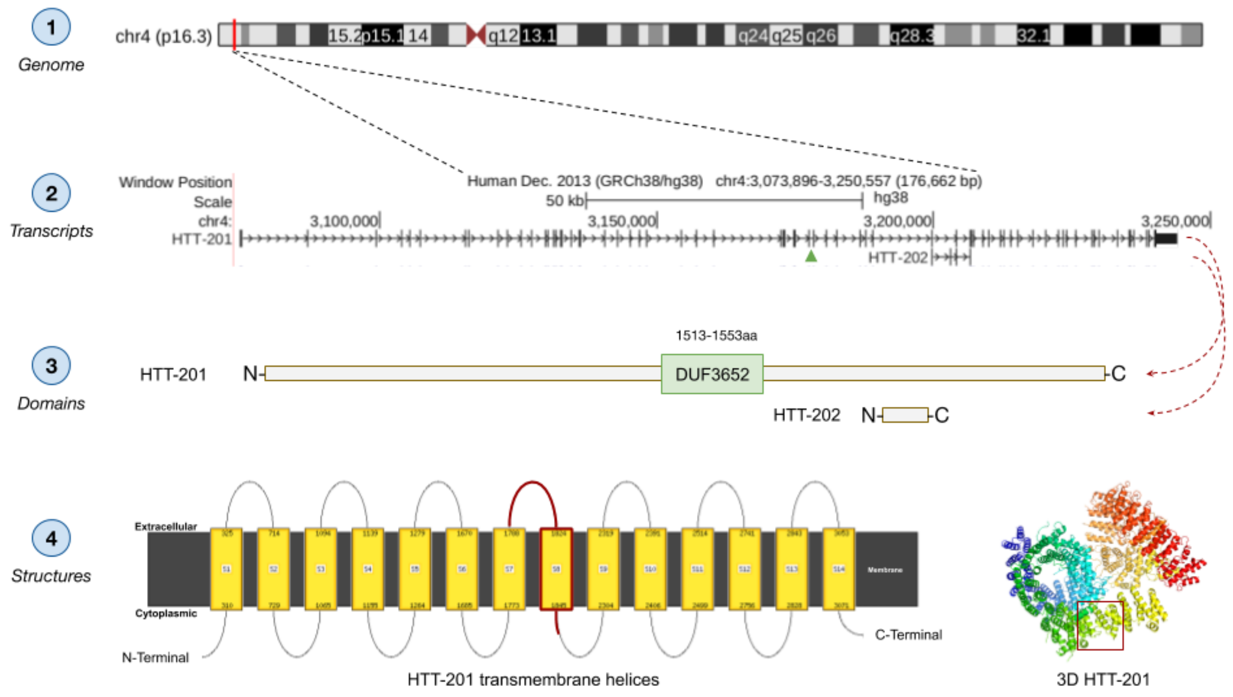


**Figure S5: Transcripts of the Huntingtin (HTT) gene.** (1) Chromosome location of HTT, which is part of the final signature that produced a score significant for survival patients with PAAD. (2) Transcripts 201 and 202 of HTT pinpointing the lack of the DUF3652 domain in the much smaller HTT-202 isoform, which is present in the final signature. (3) Transcripts 201 and 202 of HTT evidencing the position of the DUF3652 domain in the canonical HTT-201 isoform. (4) Predicted structure of HTT-201 isoform with its 14 transmembrane helices (left), highlighting in red the portion representative of HTT-202. The modeled 3D structure of HTT-201 (right) is shown in cartoon representation with rainbow coloring, where α-helix secondary structures are the helices and β-sheet secondary structures are the arrows heading to the C-terminal. The portion inside the red box is the part of HTT-201 representative of HTT-202.

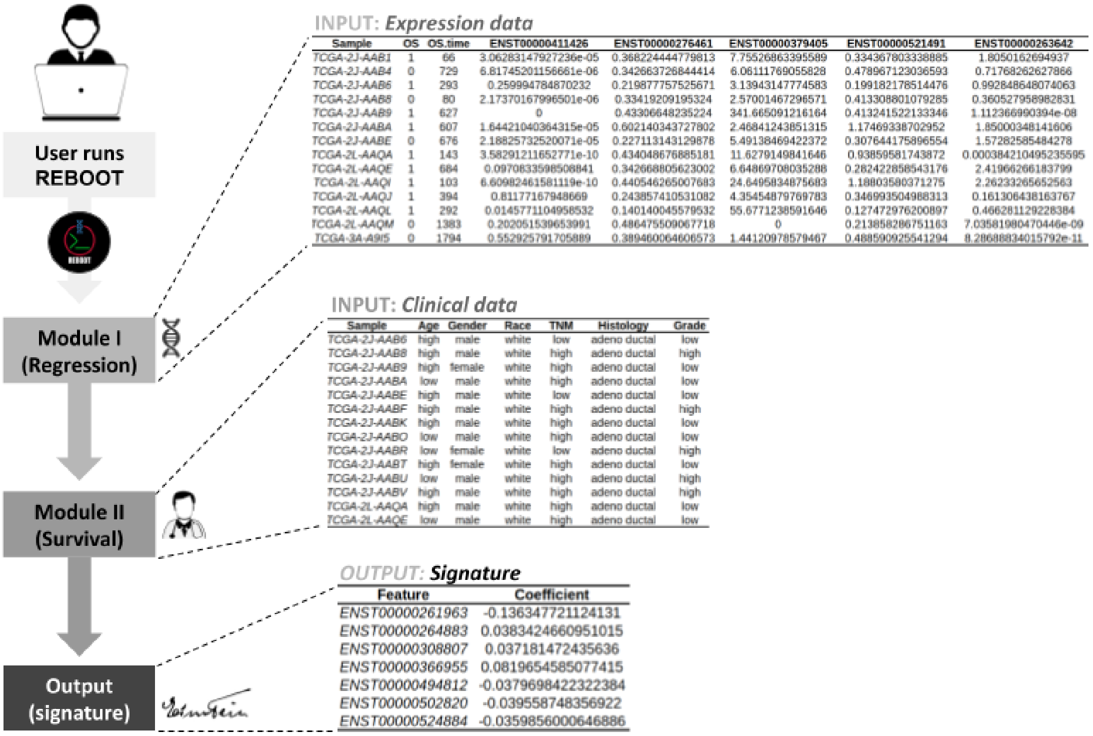


**Figure S6: Workflow of inputs to Reboot.** "Module I" requires expression data with values in TPM or FPKM. The first three columns must be the sample identifiers, patients' status (alive = 0 or dead = 1) and follow-up times (in days). "Module II" requires clinical data for multivariate analyses, where the sample identifiers match the ones present in expression data. Finally, Reboot produces several textual and graphical outputs such as a genetic signature, which is automatically used to generate a risk score evaluated in survival analyses when the "complete" mode is chosen.
